# CryoJAM: Automating Protein Homolog Fitting in Medium Resolution Cryo-EM Density Maps

**DOI:** 10.1101/2024.07.10.602952

**Authors:** Jackson Carrion, Mrunali Manjrekar, Anna Mikulevica

## Abstract

Obtaining atomic structures of large protein complexes from medium-resolution cryogenic electron-microscopy (cryo-EM) density maps is a critical bottleneck in the cryo-EM workflow. CryoJAM aims to automate this process by using a 3D Convolutional Neural Network model within a U-Net architecture. This model is trained on a novel loss function that leverages Fourier-Shell Correlation (FSC), as a proxy for quality of fit, and Root Mean Squared Error (RMSE) to help optimize fits within real space. Capitalizing on the gold-standard status of FSC in cryo-EM, this method introduces an innovative implementation of FSC into cryo-EM model fitting software, enhancing the precision and efficiency of structural analysis. After 25 epochs, CryoJAM successfully reduced the RMSE in 21 out of 26 of the test cases, effectively fitting homologous protein structures into medium-resolution cryo-EM densities.

## 1. Introduction

Cryogenic electron microscopy (cryo-EM) has revolutionized structural biology by enabling the visualization of protein complexes at atomic detail. This technique allows for researchers to study large biomolecular assemblies that are often inaccessible through other experimental or predictive methods. (Hanske et al., 2018) Structural biologists typically use high-resolution maps (sub-3Å) afforded by cryo-EM to refine the atomic structure of initial models, often derived from known protein homologs. (Liebschner et al., 2019) However, when the density map is less refined and above 3Å, this model fitting step requires considerable expert feedback, is time-consuming, and constitutes a significant bottleneck in almost every cryo-EM workflow.

In the typical workflow, the reconstruction step uses the gold standard Fourier shell correlation (FSC) to assess the quality of cryo-EM density maps. (Scheres & Chen, 2012) However, with current model fitting methods, including those that employ molecular dynamics flexible fitting (MDFF) (Singharoy et al., 2016)(Kim et al., 2019) or integrate predictions from advanced protein structure prediction models like AlphaFold (Shor & Schneidman-Duhovny, 2024) (Jamali et al., 2024) (Wang et al., 2024), are computationally expensive and often impractical for routine use on large complexes above 500kDa and at resolutions greater than 4Å . These methods struggle with the size and complexity of mega-Dalton complexes, where multiple subunits and potential conformational heterogeneity complicate straightforward predictions.

In response, there is a pressing need for innovative computational strategies that can efficiently and accurately integrate atomic-scale structural data into large medium-resolution density maps without intensive manual oversight. Here, we introduce a deep learning model, CryoJAM, a 3D Convolutional Neural Network (CNN) designed with a composite loss function that incorporates Fourier-Shell Correlation (FSC) as a proxy for quality of fitting. This model is aimed to automate and enhance the fitting of protein homologs into electron microscopy (EM) density maps. This tool accelerates the modeling of large protein complexes, reduces the need for high-resolution maps and intensive manual curation, and overall enhances the accuracy and feasibility of large-scale structural investigations in the biological sciences.

## 2. Methods

This research utilizes advanced deep learning architectures to refine protein structures by fitting them into medium-resolution cryo-EM density maps. Specifically, given a set of atomic coordinates, *C*_*h*_, representing a homolog of the target protein, *C*_*p*_, CryoJAM will generate adjusted coordinates, 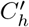, that are consistent with *C*_*p*_ and the corresponding experimentally determined medium-resolution volume *V*_*p*_. Implementing a 3D-CNN/U-Net architectures, enhanced with a novel loss function that combines elements of FSC and RMSE, will ensure both high accuracy and structural fidelity. Our generative process will involve the intermediate production of 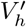to enable using FSC as a loss function.

### 2.1 Dataset

We utilize a previously published, curated dataset (He et al., 2022), comprising 256 non-redundant multi-chain protein complexes and their experimental volumes with resolutions ranging from 4Å to 8Å . To simulate greater conformational variation during fitting, synthetic homologs were generated. Each PDB structure was chosen, and one of its available chains was randomly selected. These selected chains were then subjected to rotations between 3 and 7 degrees, on a randomly chosen axis of rotation. Gaussian smoothing was then applied over *C*_*α*_ atoms to generate synthetic EM densities.

### 2.2. Data Preprocessing

To alleviate significant pose estimation challenges and highlight heterogeneous subunits, training data structures and ground truth structures were aligned based on the protein backbone as seen in Figure 1. To further reduce computational cost further, all 3D volumes (EM density maps and PDB structures) were standardized into 64^3^ voxel arrays.

**Figure 1.**
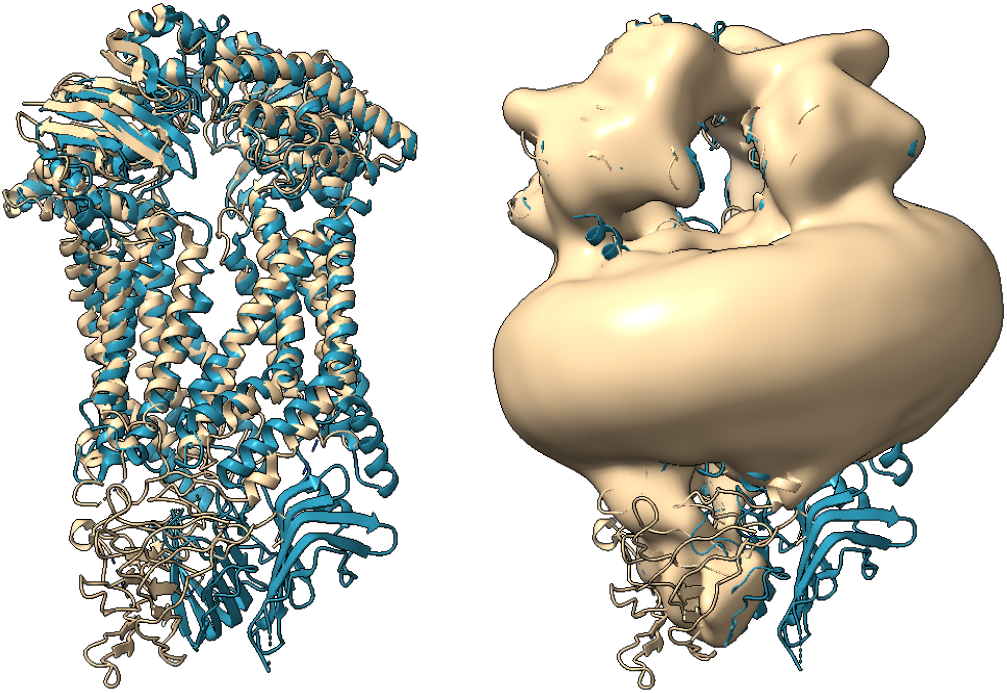
The tan atomic structure (left) represents the true cryo-EM derived *Homo sapiens* LptB2FG Transporter protein, which serves as the ground truth in this study. The corresponding medium-resolution density map is shown on the right. This density map was experimentally determined and serves as the target for fitting. The blue atomic structure illustrates the homologous *E. coli* LptB2FG Transporter protein, derived from X-ray crystallography, which requires fitting into the EM density map.

### 2.3. Model Approach

In our study, CryoJAM is a multi-modal UNet-based architecture tailored for 3D volumetric data analysis, designed specifically to handle both EM densities and homolog structures as 3-dimensional tensors. These tensors are packaged together into a final input tensor of shape [1, 2, 64, 64, 64], with the batch size constrained to one. This design facilitates simultaneous processing of stacked homolog atomic structures and EM density maps. The output of the model is a [1, 1, 64, 64, 64] tensor that represents the adjusted homolog coordinates as a volume of continuous values from 0 to 1. This method enables the computation of both the RMSE as well as the FSC between the predicted structure and the true atomic coordinates, which also are also pre-processed and repressed as 64^3^ voxel arrays. With 22,464,223 trainable parameters, CryoJAM achieves precise adjustments that ensure an optimal fit between predicted structures and EM density maps. The model’s performance, validated across 26 diverse test cases, demonstrates its effectiveness. For a detailed explanation of the algorithmic framework behind CryoJAM, refer to Appendix A.

#### 2.3.1. Composite Loss Function

While FSC provides a quantitative measure of the similarity between the reconstructed and target density volumes via Fourier space, RMSE offers a more direct comparison of atomic accuracy. Note that a coordinate based RMSD would require matching between atoms in the predicted and ground-truth. And such matching from a volume would require a heuristic that would complicate differentiation and backpropagation. Therefore, our model’s training focuses on RMSE and FSC losses. For a detailed explination of the FSC and RMSE calculations please refer to Appendix A.

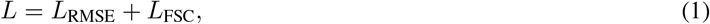

where:

- *L*_RMSE_ represents the loss component corresponding to the RMSE, aiming for minimization.
- *L*_FSC_ corresponds to the FSC metric, treated here as 1 - FSC, to facilitate maximization within a minimization framework.

The RMSE component is defined as:

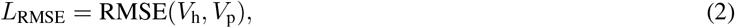

where *V*_h_ and *V*_p_ are the predicted and ground truth protein volumes, respectively.

The FSC component is defined as:

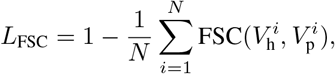

where 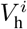and 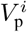are the predicted and ground truth volumes in the *i*-th shell of the Fourier space, and *N* is the total number of shells.

### 2.4. PDB Generation

Our model generates a volume reflecting its predictions of Carbon alpha (*C*_*α*_) atoms within the EM density in a way that preserves volumetric spatial congruence while retaining the most intense pixels as the backbone density. However, because the volumes generated are composed of continuous scalars, we developed a small post-processing workflow to generate realistic all-atom coordinates from the given volume. First, given *K* number of *C*_*α*_ atoms in the structure, a KD-tree (Skrodzki, 2019) is used to pick out the top *K* voxel activations from the volume subject to a minimum distance of 1Å, to binarize the generated volume and output a set of 3D coordinates sufficient to build a PDB file. After reverting the structure-specific unscaling normalization applied during pre-preprocessing, a greedy matching algorithm is then used to help match these selected *C*_*α*_ atoms to their closest atom in the input structure to adjust accordingly. These new *C*_*α*_ traces are then processed using PULCHRA (Rotkiewicz & Skolnick, 2008)(Terashi et al., 2024), which can rapidly construct physically realistic all-atom structures given only *C*_*α*_ coordinates, enabling the visualization of all-atom structure in tenths of a second. Taken together, an inference and post-processing pass for each prediction takes about 9 seconds, with the bulk of time being attributed to the greedy assignment.

### 2.5. Validation

Validation of CryoJAM involves the use of multiple metrics, including FSC and RMSE. Low RMSE values are particularly crucial as it indicates a high degree of accuracy in the 3D alignment between the predicted structure and the true structure. This metric can ensure that CryoJAM predicts reliable structural interpretations. To provide holistic verification of the fitting accuracy, we also utilized advanced visualization tools such as ChimeraX. (Goddard et al., 2018) These tools allowed us to visually inspect and compare the adjusted protein structures within the EM density maps.

## 3. Results

CryoJAM yields compelling results validating the efficacy of deep learning models when tasked with fitting atomic models into medium resolution density maps. Predictions after only 25 epochs (approximately 2 min per epoch) demonstrated the model was successful in improving the FSC and RMSE metric for 22 out of 26 of the test cases and matching MD simulation quality of fitting. Further comparisons even show 69% of the test cases achieved an RMSE value less than or equal to 1 when compared to the ground truth structure.

### 3.1. Improving FSC and RMSE

The model’s learning trajectory is apparent in the convergence of the composite loss function curves seen in Figure 2. Furthermore, FSC and RMSE analyses depict consistent reductions for 22/26 and 21/26 test cases respectively as seen in Figure 3. Notably, 69% of the test cases achieved an RMSE value less than or equal to 1, demonstrating the model’s ability to generate highly accurate fits. Figure 4 further exemplifies the improvements seen in the FSC metric for a specific test case, as we see from the maximized FSC curve after model processing, illustrating the prediction is much closer to ground truth. For a comprehensive analysis of the model’s performance across the 26 test cases, refer to Appendix B.

**Figure 2.**
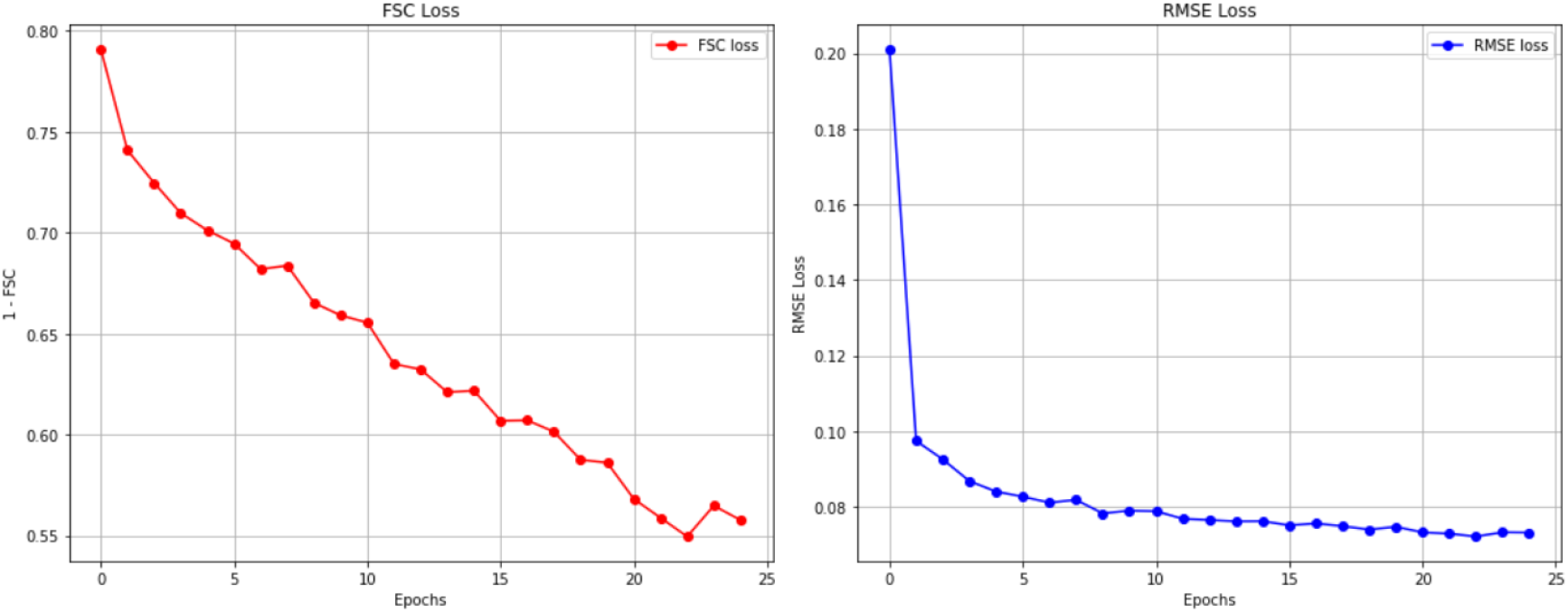
Depiction of the individual loss curves for FSC loss, expressed as 1 - FSC, and the RMSE loss across training epochs. The left panel shows the FSC loss curve, indicating a steady decrease as the model trains, reflecting an increasing similarity between the predicted and target density volumes in Fourier space. The right panel shows the RMSE loss curve, which rapidly converges during the early epochs, indicating that the model quickly learns to minimize positional discrepancies between predicted and actual atomic positions in real space.

**Figure 3.**
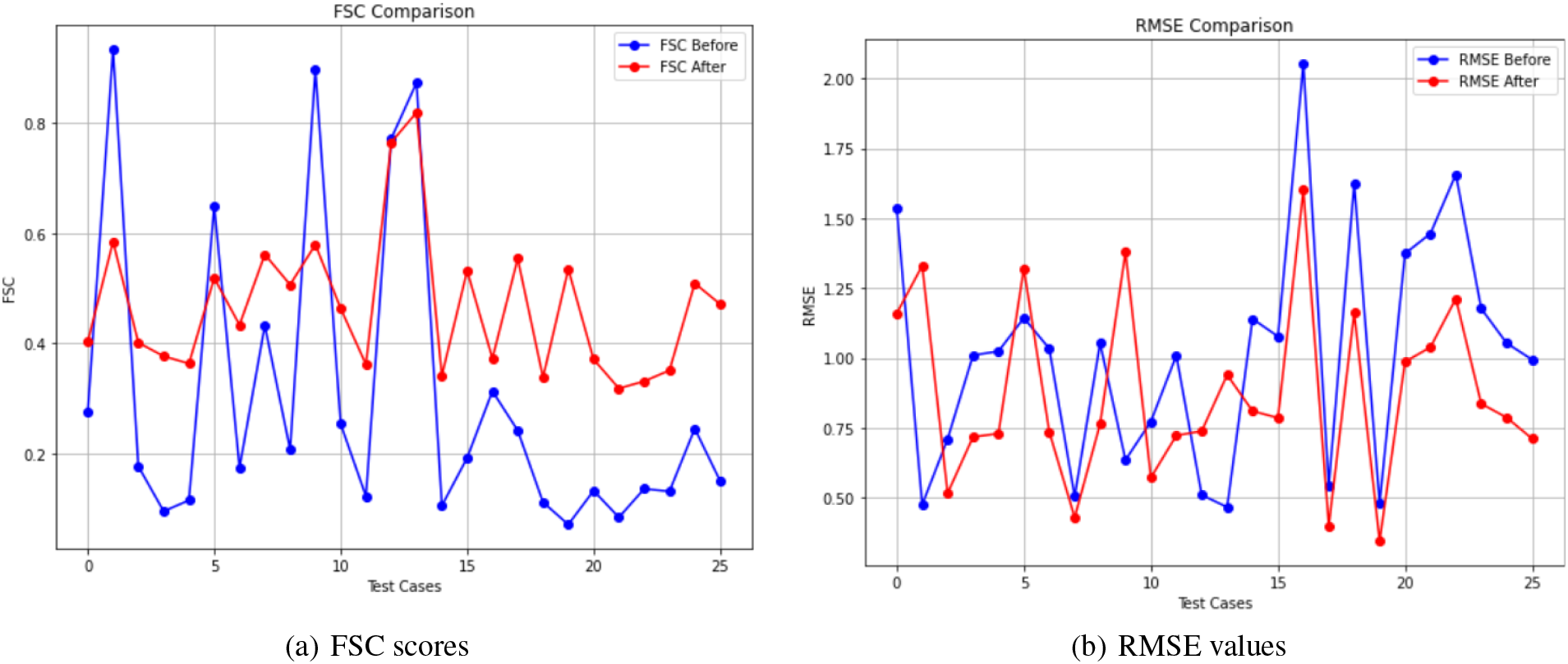
A comparison of FSC scores (a) and RMSE values (b) between homolog structure and predicted structure, compared with the ground truth, across 26 test cases. (a) The blue line represents FSC scores for homolog structure, while the red line depicts FSC scores for predicted structure. (b) The blue line represents RMSE values for the protein backbone before our model’s adjustment, while the red line depicts RMSE values after adjustment.View table in Appendix B.

**Figure 4.**
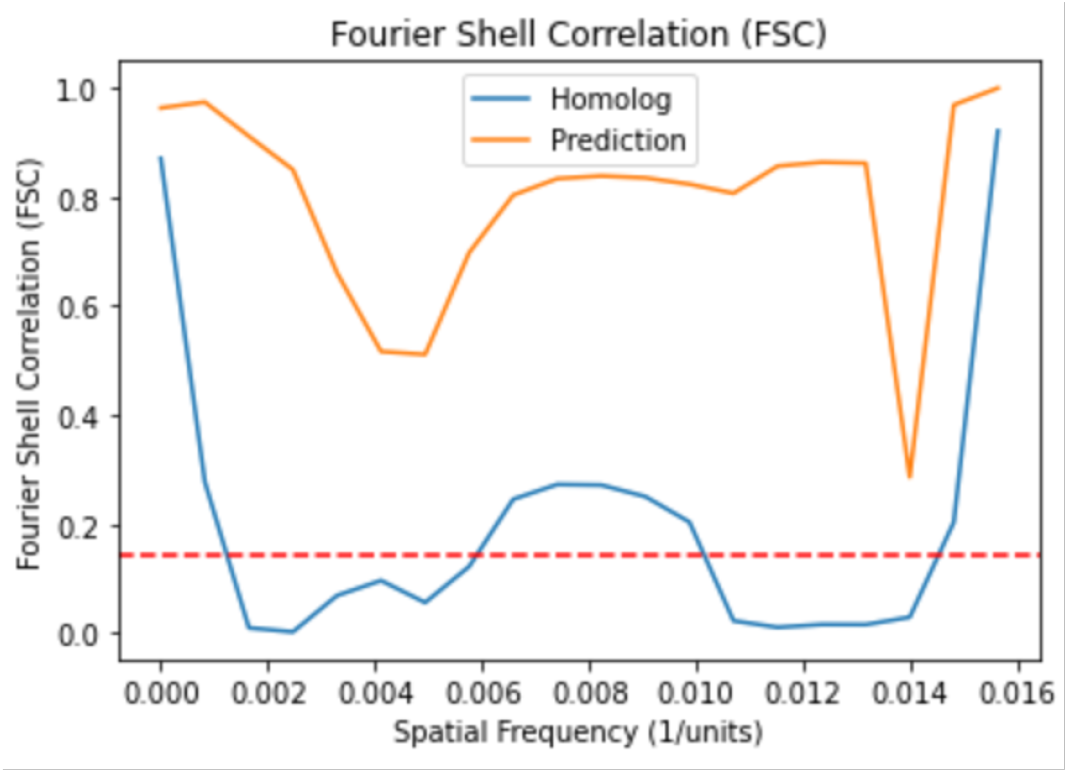
The FSC curve that shows the comparison between ground truth and homolog structure and ground truth and the predicted structure. The blue line represents the FSC between the true EM density and the homolog/input structure, while the orange line represents the FSC between the true EM density and the predicted structure generated by CryoJAM. The FSC curve illustrates the spatial frequency correlation between the volumes. Higher FSC values, as seen in the orange line, indicates a closer match to the true EM density. The dashed red line represents the 0.143 threshold, a commonly used criterion for assessing the resolution and quality of two EM density maps.

## 4. Discussion

The results of our study underscore the efficacy of CryoJAM in automatting the fitting of known protein structures into medium-resolution cryo-EM density maps. The models dual optimization strategy, leveraging both FSC and RMSE proved effective in achieving high-fidelity fits.

### 4.1. Analysis of the Loss Function

The analysis of the composite loss function, Figure 2, reveals insight into the model’s learning process. RMSE, which relies on real space calculations, is predominantly learned during the earlier phases of training. While the FSC loss is minimized more gradually, reflecting the model’s ongoing refinement of high-frequency details and volumetric consistency. Together, these metrics demonstrate complementary contributions to the model’s learning process, with each targeting distinct aspects of the structural prediction task.

As the two major goals of the loss function was to minimize FSC loss (1-FSC) and RMSE, Figure 3, as well as Appendix B, illustrates the consistent increase in performance (higher FSC and lower RMSE) for 81% of the test cases. Figure 3(a) depicts the predicted test structures consistently showing significantly higher FSC scores compared to ground truth in 22 out of 26 test cases. Likewise, Figure 3(b) and the comprehensive table in Appendix B. highlights initial discrepancies in RMSE levels, with the model consistently yielding significantly lower RMSE scores compared to the ground truth in 22 out of 26 test cases. 18 out of the 26 predicted structures also achieved an RMSE less than or equal to 1, suggesting the predicted structure aligns significantly well with the true structure.

### 4.2. Ablation Study of the Loss Function

To further validate and explore the effects of each component in the composite loss function, an ablation study was included in Appendix B. From these results, we see for a number of test examples, incorporating FSC into the loss function alongside RMSE improves fitting quality. The addition of FSC into the loss function posed advantageous to reducing overfitting while training.

### 4.3. Limitations

Of the 6 test cases where the FSC was not increased, 3 out of the 6 cases had very similar structures compared to the ground truth, suggesting that our model may struggle more with determining small differences between very similar homolog structures. Low-resolution EM density, resolutions exceeding 7Å and those containing membrane/detergent density, also present notable challenges. Our model exhibited lower FSC scores for predicted structures within such densities, ultimately suggesting these noisy FSC shells that correlate with detergent micelles may hinder our model’s ability to accurately predict all classes of proteins despite a diverse training set.

### 4.4. Benchmarking

Compared to physics-based methods like MD simulations, as seen in Figure 5, we can visualize our predicted protein model also achieving extremely similar positioning without the need for costly simulations. In Figure 5, we can see the teal (predicted) structure are not only resident within the density map, but also overlapping with a majority of the physics-based structure seen in orange. This orange structure was manually created and refined with MD simulations as seen in previous studies (Snijder et al., 2017). The right-most alpha helix in Figure 5 displays a more difficult adjustment task, as the tan structure is already within the density. This highlights CryoJAM’s ability to accurately identify and adjust specific backbone atoms within the protein to achieve an optimal fit in the medium resolution map. Nonetheless, with our model capable of making new predictions in the order of seconds, this shows the significance in a deep learning approach compared to time-consuming simulations.

**Figure 5.**
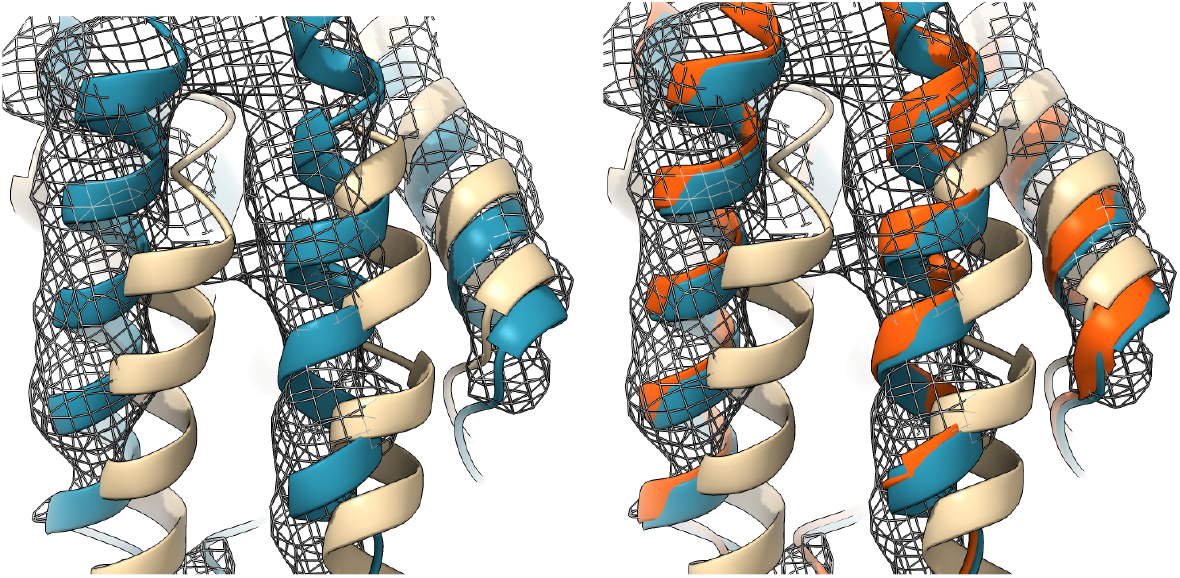
The image showcases three protein structures within an electron density map mesh of 5N8Y. The predicted structure (depicted in blue) exhibits a precise alignment with the EM density mesh and the manual and MD-based ground truth structure (depicted in orange), contrasting with the input homologous structure (depicted in tan).

## 5. Conclusions

CryoJAM is a 3D CNN architecture tailored specifically for the automation of model fitting in the cryo-EM workflow. This deep learning architecture leverages a novel composite loss function that combines Fourier-Shell Correlation and Root-Mean-Square Error to effectively proxy the quality of fitting into medium resolution density maps. The FSC is an integral component of our model, allowing for accurate yet quick calculations to drive the fitting process. Our results demonstrate the model’s robust performance, achieving lower RMSE values in 81% of test cases and shows accurate fitting when compared to ground truth structures; underscoring the model’s efficiency and highlighting its potential to streamline the cryo-EM workflow. By automating the model fitting process and validating new proxies and loss functions, CryoJAM can significantly accelerate cryo-EM structural determinations, making challenging proteins targets more accessible to atomic-level observations.

## Software and Data Availability

Training was conducted on a NVIDIA-A10 GPU for 25 epochs. Our proposed UNet architecture consists of 22,464,223 total trainable parameters and requires 10.2 GB of GPU memory. All code can be found on our GitHub. https://github.com/jtcarrion/CryoJAM.

## Acknowledgments

We thank Joey Davis and Sergey Ovchinnikov for their valuable feedback throughout this project and for providing access to essential computing resources. Their support and guidance were crucial to the successful completion of this work.

## A. Appendix

Here we can see the underlying algorithm behind the code

### Algorithm

Protein Fitting

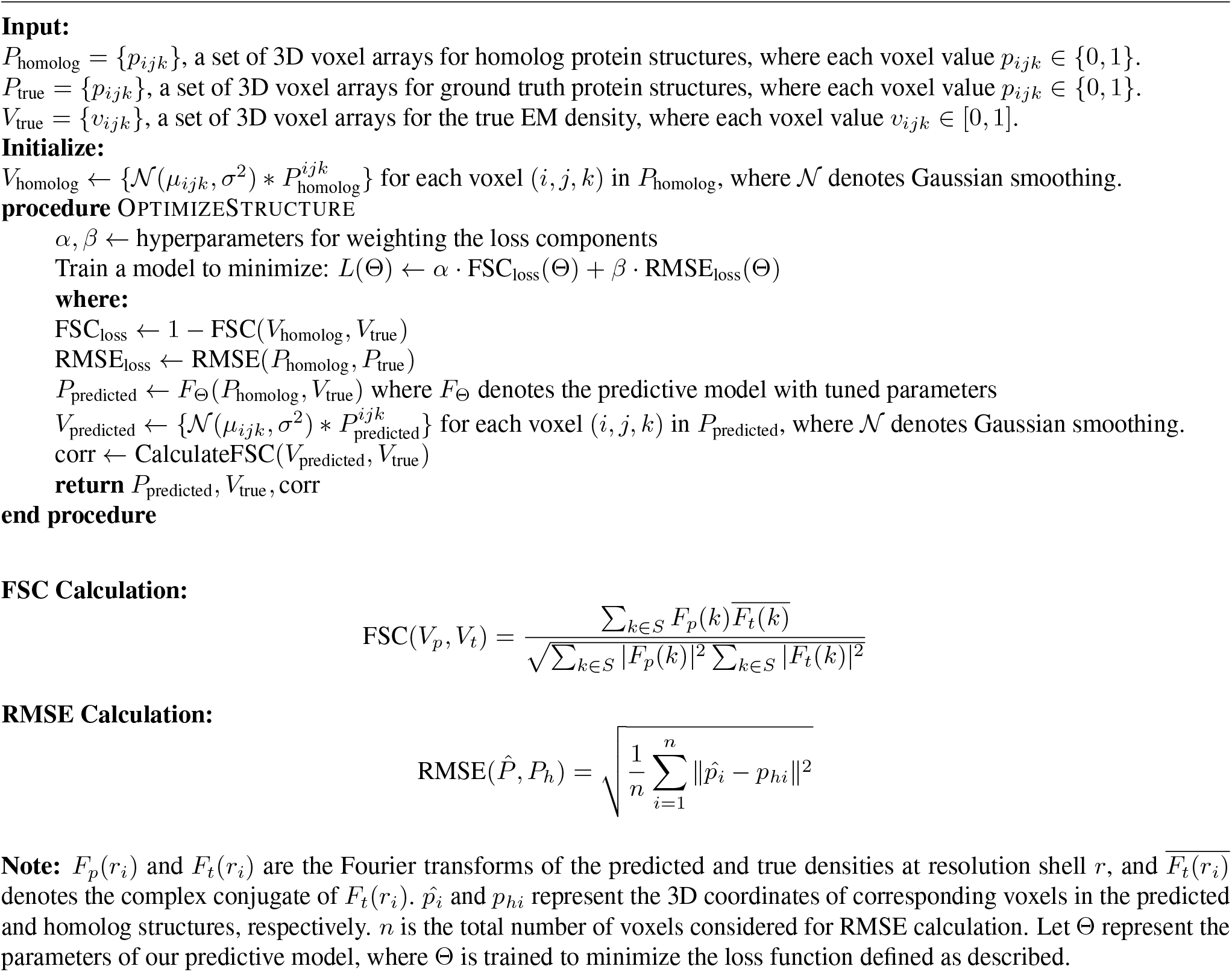

## B. Appendix

Tabel below shows the RMSE values for 26 test cases pre and post-fit compared to the ground truth structure. Also shows results from the ablation study on the loss function suggesting that a composite loss function reduced over-fitting and improved results.

| PDB ID | RMSE Before | RMSE After | RMSE After (w/o FSC) |
| --- | --- | --- | --- |
| 5W1R | 1.534 | 1.159 | 1.13 |
| 5G5L | 0.476 | 1.328 | 1.156 |
| 6NI2 | 0.711 | 1.388 | 0.53 |
| **6M5V** | **1.01** | **0.719** | **0.728** |
| **6PEK** | **1.024** | **0.731** | **0.741** |
| **7JPU** | **1.144** | **1.019** | **1.159** |
| 7KAL | 1.034 | 0.737 | 0.734 |
| 6SHS | 0.506 | 0.435 | 0.433 |
| **6S2E** | **1.036** | **0.671** | **0.76** |
| 3JC5 | 0.694 | 1.378 | 1.224 |
| **6C14** | **0.774** | **0.506** | **0.59** |
| 6OCE | 0.652 | 0.742 | 0.727 |
| 6RGL | 0.511 | 0.74 | 0.588 |
| 7JTH | 0.468 | 0.793 | 0.726 |
| 5FWP | 1.032 | 1.205 | 0.813 |
| 6F2D | 1.079 | 0.787 | 0.774 |
| **6EL1** | **1.064** | **0.787** | **1.531** |
| 6WYK | 1.025 | 1.068 | 0.423 |
| **6V3G** | **1.624** | **1.106** | **1.154** |
| 6MLU | 1.044 | 1.041 | 0.400 |
| **6DVW** | **1.374** | **0.897** | **0.99** |
| **7IWI** | **1.104** | **0.737** | **1.008** |
| **6LQI** | **1.658** | **0.986** | **1.174** |
| **6J1J** | **1.178** | **0.789** | **0.862** |
| 5N9Y | 1.105 | 0.884 | 0.773 |
| **6Y5K** | **1.053** | **0.713** | **0.715** |
| Avg. | 0.996 | 0.897 | 0.840 |

**Note:** The bold rows highlight the 12 test cases that resulted in a lower RMSE post-fitting when comparing composite loss function to the same UNet architecture trained with only RMSE.

## References

Goddard, T. D., Huang, C. C., Meng, E. C., Pettersen, E. F., Couch, G. S., Morris, J. H., and Ferrin, T. E. Ucsf chimerax: Meeting modern challenges in visualization and analysis. Protein Sci., 27(1):14–25, January 2018.

Hanske, J., Sadian, Y., and Müller, C. The cryo-em resolution revolution and transcription complexes. Current Opinion in Structural Biology, 2018.

He, J., Lin, P., Chen, J., Cao, H., and Huang, S. Model building of protein complexes from intermediate-resolution cryo-em maps with deep learning-guided automatic assembly. Nature Communications, pp. 4066, 2022.

Jamali, K., Käll, L., Zhang, R., Brown, A., Kimanius, D., and Scheres, S. H. W. Automated model building and protein identification in cryo-em maps. Nature, 628(8007):450–457, April 2024.

Kim, D. N., Moriarty, N. W., Kirmizialtin, S., Afonine, P. V., Poon, B., Sobolev, O. V., Adams, P. D., and Sanbonmatsu, K. Y. Cryo fit: Democratization of flexible fitting for cryo-em. Journal of Structural Biology, 208(1):1–6, 2019. doi: 10.1016/j.jsb.2019.05.012.

Liebschner, D., Afonine, P. V., Baker, M. L., Bunkóczki, G., Chen, V. B., Croll, T. I., Hintze, B., Hung, L.-W., Jain, S., McCoy, A. J., Moriarty, N. W., Oeffner, R. D., Poon, B. K., Prisant, M. G., Read, R. J., Richardson, J. S., Richardson, D. C., Sammito, M. D., Sobolev, O. V., Stockwell, D. H., Terwilliger, T. C., Urzhumtsev, A. G., Videau, L. L., Williams, C. J., and Adams, P. D. Macromolecular structure determination using x-rays, neutrons and electrons: recent developments in phenix. Acta Crystallographica Section D, 75(10):861–877, October 2019. doi: 10.1107/S2059798319011471.

Rotkiewicz, P. and Skolnick, J. Fast procedure for reconstruction of full-atom protein models from reduced representations. J. Comput. Chem., 29(9):1460–1465, July 2008.

Scheres, S. and Chen, S. Prevention of overfitting in cryo-em structure determination. Nature Methods, 2012.

Shor, B. and Schneidman-Duhovny, D. Combfold: predicting structures of large protein assemblies using a combinatorial assembly algorithm and alphafold2. Nature, 2024.

Singharoy, A., Teo, I., McGreevy, R., Stone, J. E., Zhao, J., and Schulten, K. Molecular dynamics-based refinement and validation for sub-5 Å cryo-electron microscopy maps. eLife, pp. e16105, 2016.

Skrodzki, M. The k-d tree data structure and a proof for neighborhood computation in expected logarithmic time. CoRR, abs/1903.04936, 2019.

Snijder, J., Schuller, J. M., Wiegard, A., Lössl, P., Schmelling, N., Axmann, I. M., Plitzko, J. M., Förster, F., and Heck, A. J. R. Structures of the cyanobacterial circadian oscillator frozen in a fully assembled state. Science, 2017.

Terashi, G., Wang, X., Prasad, D., and Kihara, D. Deepmainmast: integrated protocol of protein structure modeling for cryo-em with deep learning and structure prediction. Nat Methods, 2024.

Wang, X., Zhu, H., Terashi, G., Taluja, M., and Kihara, D. Diffmodeler: Large macromolecular structure modeling in low-resolution cryo-em maps using diffusion model. bioRxiv, 2024.

